# Shiny AMMOA: an interactive platform for integrative multi-omics analysis of murine aging

**DOI:** 10.64898/2026.05.18.726091

**Authors:** Mayuka Ninomiya Kanda

**Affiliations:** Department of Bioengineering and QB3 Institute, University of California, Berkeley, Berkeley, CA 94720, USA

**Author notes:** Corresponding Author Mayuka Ninomiya Kanda.

**Keywords:** Multi-omics, Transcriptomics, Proteomics, Metabolomics, R Shiny, Bioinformatics

## Abstract

Aging is accompanied by complex, tissue-specific molecular changes across multiple biological layers, yet integrative analysis of multi-omics datasets remains challenging for many experimental researchers due to technical and computational barriers. Here, I present Shiny Aging Murine Multi-Omic Analyzer (Shiny AMMOA), a graphical user interface (GUI)-based, user-friendly analytical platform that enables interactive exploration of murine aging-associated bulk transcriptomic, proteomic, and metabolomic datasets. Shiny AMMOA integrates publicly available multi-omics resources within a unified R Shiny framework and supports end-to-end analyses, including differential expression testing, pathway enrichment analysis, and pathway-level visualization across individual and multiple omics layers.

Using representative use cases, I demonstrate that Shiny AMMOA recapitulates key findings from original source studies and facilitates intuitive discovery of tissue-, pathway-, and modality-specific aging signatures, including age-associated alterations in unfolded protein response, extracellular matrix organization, and metabolic pathways across specific tissues and omics layers. The platform further enables integrated visualization of molecular changes across omics layers on Kyoto Encyclopedia of Genes and Genomes (KEGG) pathway diagrams, supporting hypothesis generation at the systems level.

By democratizing access to integrative multi-omics analysis while preserving analytical rigor, Shiny AMMOA provides an extensible resource for experimental biologists and aging researchers to interrogate large-scale public datasets, prioritize biological pathways, and accelerate translation of multi-omics insights into testable experimental hypotheses. Shiny AMMOA is available at https://github.com/M-Ninomiya-Kanda/Shiny_AMMOA_local, and a lightweight web-based demonstration version with limited functionality is available at https://m-ninomiya-kanda.shinyapps.io/shiny_ammoa_web/.

## Background

Aging is a major risk factor for numerous chronic diseases, including cardiovascular, neurodegenerative, and metabolic disorders, and ultimately contributes to increased mortality [1,2]. Accordingly, understanding biomolecular changes associated with aging is of critical importance for identifying potential therapeutic strategies to promote healthy aging. Because aging affects multiple biological layers simultaneously [3,4], comprehensive characterization of age-associated molecular alterations often requires integrative analyses across multiple omics layers.

With advances in high-throughput technologies, non-targeted exploratory analyses of genome, epigenome, RNA expression, protein abundance, and metabolite levels, or collectively referred to as multi-omics, are increasingly applied in aging research. However, generating multi-omic datasets from scratch requires substantial technical expertise, financial resources, and, in preclinical studies, large numbers of laboratory animals. To mitigate these costs, reduce redundant animal use for ethical considerations, and promote open science, it has become standard practice to deposit raw omics data in public repositories upon publication [5]. However, such datasets often remain difficult to reuse despite their availability, as data accessibility and preprocessing levels vary widely across studies. In many cases, ready-to-analyze, curated datasets are not provided, creating a barrier for experimental scientists who may not be familiar with advanced statistical analyses.

Some studies have addressed this challenge by offering interactive, web-based data browsers that allow users to explore the omics datasets generated in their original studies [6–8]. While these tools facilitate intuitive exploration of individual expression patterns, they are typically designed for visualization purposes only and do not support advanced biostatistical analyses such as pathway enrichment or network-based investigation. Conversely, several graphical user interface (GUI)-based applications have been developed to enable sophisticated statistical analyses and visualization of omics data [9–11]. However, these tools generally require users to upload their own curated datasets, placing another burden of data preprocessing and quality control on the users’ end and creating an additional hurdle for non-coding experimental scientists. Together, these limitations highlight the need for an end-to-end, GUI-based platform that integrates data retrieval, preprocessing, statistical analysis, and visualization.

At the level of biological interpretation, pathway enrichment analysis is a widely used approach for interpreting non-targeted omics datasets, including those generated in aging studies [12]. Although enrichment results can provide valuable insights into biological processes affected by aging, they often do not reveal which specific molecules within a pathway drive the observed changes. Canonical pathway annotations typically encompass diverse functional components, such as extracellular signaling molecules, membrane receptors, cytosolic signaling mediators, and nuclear transcription factors, without explicitly accounting for their distinct roles or spatial organization [13,14]. Visualizing expression changes of all pathway components directly on pathway diagrams therefore represents an effective strategy for gaining more intuitive and detailed mechanistic insight into molecular alterations [15].

Here, I present Shiny Aging Murine Multi-Omic Analyzer (Shiny AMMOA), a GUI-based application developed using the R Shiny framework. Shiny AMMOA enables exploratory analysis of age-associated molecular changes across three omics layers (bulk RNA-seq, proteomics, and metabolomics) across multiple tissues in *Mus musculus* from publicly available sources. By combining differential expression analysis, pathway enrichment analysis and pathway-level visualization of gene, protein, and metabolite changes, AMMOA provides an accessible platform for experimental scientists to investigate publicly available aging-related multi-omics datasets without requiring extensive computational expertise.

## Methods

### Data Source

All source data used in Shiny AMMOA, including bulk RNA-seq [6], proteome [16] and metabolome [7], were retrieved from publicly available published sources and downloaded from the data repository indicated in the original publications. Specifically, bulk RNA-seq data were obtained from the Gene Expression Omnibus (GEO) under accession number GSE132040, proteomic data were retrieved from FigShare (DOI: 10.6084/m9.figshare.19765849), and metabolomic data were extracted from the Supplementary Table of the original publication. Further details of each dataset are provided in the corresponding original studies. Briefly, the bulk transcriptomic dataset was derived from the Tabula Muris Senis bulk RNA-seq resource, a comprehensive murine aging atlas spanning multiple tissues and age groups. Bulk RNA-seq profiles were obtained from male and female C57BL/6JN mice across early adulthood to advanced age, with samples collected from 17 distinct tissues. Proteomic data were obtained from a discovery-based tandem mass tag (TMT) proteomics study characterizing age- and sex-associated proteomic changes across murine tissues. This dataset comprises quantitative proteome measurements from 10 tissues collected from C57BL/6J mice at 8 and 18 months of age, including both male and female animals. Metabolomic data were sourced from a comprehensive study investigating age-related metabolic alterations in mice. The dataset includes metabolite abundance profiles from 15 tissues, including serum, obtained from young (3–6 months) and aged (>21 months) C57BL/6J mice of both sexes. Metabolites were quantified using hydrophilic interaction chromatography coupled with high-resolution quadrupole–Orbitrap mass spectrometry.

### Bulk RNA-seq Data Preprocessing

Raw bulk RNA-seq count data were downloaded from the GEO under accession number GSE132040. Using the same quality filtering criteria as applied in the original publication [6], samples with a total read count below 4 million were excluded from subsequent analyses. The raw count data matrix was analyzed using the standard DESeq2 pipeline [17], with sex and age included as covariates in the design formula. In differential expression analyses comparing two age groups (e.g., 3 months vs. 21 months), age was treated as a binary variable. In contrast, for analyses including all age groups (i.e., 1, 3, 6, 9, 12, 15, 18, 21, and 24 months), age was modeled as a continuous variable after scaling. In the all-age-group analysis, the scaling was defined such that a log2 fold change of 1 corresponds to a doubling of mean expression over a 12-month interval.

### Proteomic Data Preprocessing

For differential expression analyses of the proteomic data, intermediate data files generated by the authors of the original publication [16] were retrieved from FigShare (DOI: 10.6084/m9.figshare.19765849) and used in Shiny AMMOA. Briefly, in the original publication, protein abundance estimates were obtained following batch-aware normalization and ratio scaling and were represented on a log2 scale. Subsequently, abundance values were modeled by the original authors using ordinary least squares (OLS) regression, with age, sex, and analytical batch included as covariates. Under this modeling framework, the estimated age coefficients correspond to covariate-adjusted log2 fold changes between young (8-month-old) and aged (18-month-old) mice. Accordingly, these values are labeled as “covariate-adjusted Log2FC” in the Shiny AMMOA application.

### Metabolomic Data Preprocessing

Metabolite abundance data were obtained as processed ion count data from the supplementary data table of the original publication [7]. Differential abundance analysis was performed using OLS regression, with log2-transformed ion counts as the response variable and age and sex included as covariates. In this modeling framework, the estimated age coefficients represent covariate-adjusted log2 fold changes between young and aged mice. Unlike the original study, interaction terms were not included in the OLS model, in order to maintain consistent covariate adjustment across omics layers such as the transcriptome and proteome. To evaluate the impact of this simplification, I confirmed that my analysis produced results comparable to those reported in the original study, indicating that the omission of interaction terms had minimal effect on the overall conclusions. Metabolite common names were mapped to Kyoto Encyclopedia of Genes and Genomes (KEGG) and Human Metabolome Database (HMDB) identifiers using MetaboAnalyst 6.0 (web version) [18], with manual curation.

### Enrichment Analysis

In Shiny AMMOA, pathway enrichment analysis is performed using an over-representation analysis (ORA) framework. For transcriptomic and proteomic data, ORA querying KEGG and Gene Ontology (GO) pathways is conducted using clusterProfiler (version ≥ 4.14.6). For metabolomic data, ORA querying KEGG pathways is also performed using clusterProfiler. In contrast, ORA querying the Small Molecule Pathway Database (SMPDB) is performed using MicrobiomeProfiler (version ≥ 1.12.0), which operates as an extension of and depends on clusterProfiler. Differentially expressed genes, proteins, or metabolites are defined based on user-specified thresholds for the Benjamini–Hochberg–adjusted p-value (false discovery rate, FDR) and log2 fold change, and are subsequently submitted to the corresponding enrichment analyses. Enrichment results are visualized as dot plots. For transcriptomic and proteomic analyses, when more than 10 significant (adjusted p-value < 0.05) pathways are identified, pathways are grouped into five clusters based on the similarity of their component genes or proteins. Semantic similarity is computed using enrichplot (version ≥ 1.26.6), and clustering is performed using Ward’s D2 hierarchical clustering algorithm implemented in stats (version ≥ 4.4.2). When more than 25 significant pathways are detected, only the top five pathways from each cluster, ranked by adjusted p value, are displayed to reduce redundancy.

### Kyoto Encyclopedia of Genes and Genomes Pathway Mapping

On the “Pathway Mapping” page, age-associated changes in gene, protein, and metabolite expression are visualized using the pathview package [15] (version ≥ 1.46.0). Pathview is an R/Bioconductor package that maps user-provided expression data onto KEGG pathway diagrams using a heat color scale, enabling intuitive visualization of the locations of up- and down-regulated molecules within each pathway. In Shiny AMMOA, rectangular nodes represent genes or proteins, whereas small circular nodes represent metabolites. Differential expression results, represented as log2 fold change values, are passed to the pathview() function, and the resulting heat-colored pathway diagrams are displayed in the user interface.

### R Shiny Application Development and Implementation

Shiny AMMOA was implemented as a graphical user interface–based application using the R Shiny framework to enable interactive exploration of multi-omics data without requiring direct programming by the user. The overall structure of the application follows the schematic workflow shown in Figure 1. First, omics datasets and associated analysis results are retrieved from publicly available repositories (see the “Source Data” section) and incorporated into the application as structured data objects. Second, these datasets undergo preprocessing steps specific to each omics layer (see the corresponding data preprocessing sections in the Methods) and are stored for downstream use. Third, selected analyses are performed dynamically within the application in response to user input. Within this workflow, users specify a set of analysis parameters, such as the data type (bulk RNA-seq, proteome, or metabolome), tissue of interest, and thresholds for differential expression and pathway enrichment. Based on these inputs, the application executes on-demand analyses, including pathway enrichment and pathway visualization, and displays the results through interactive plots and tables.

**Figure 1.**
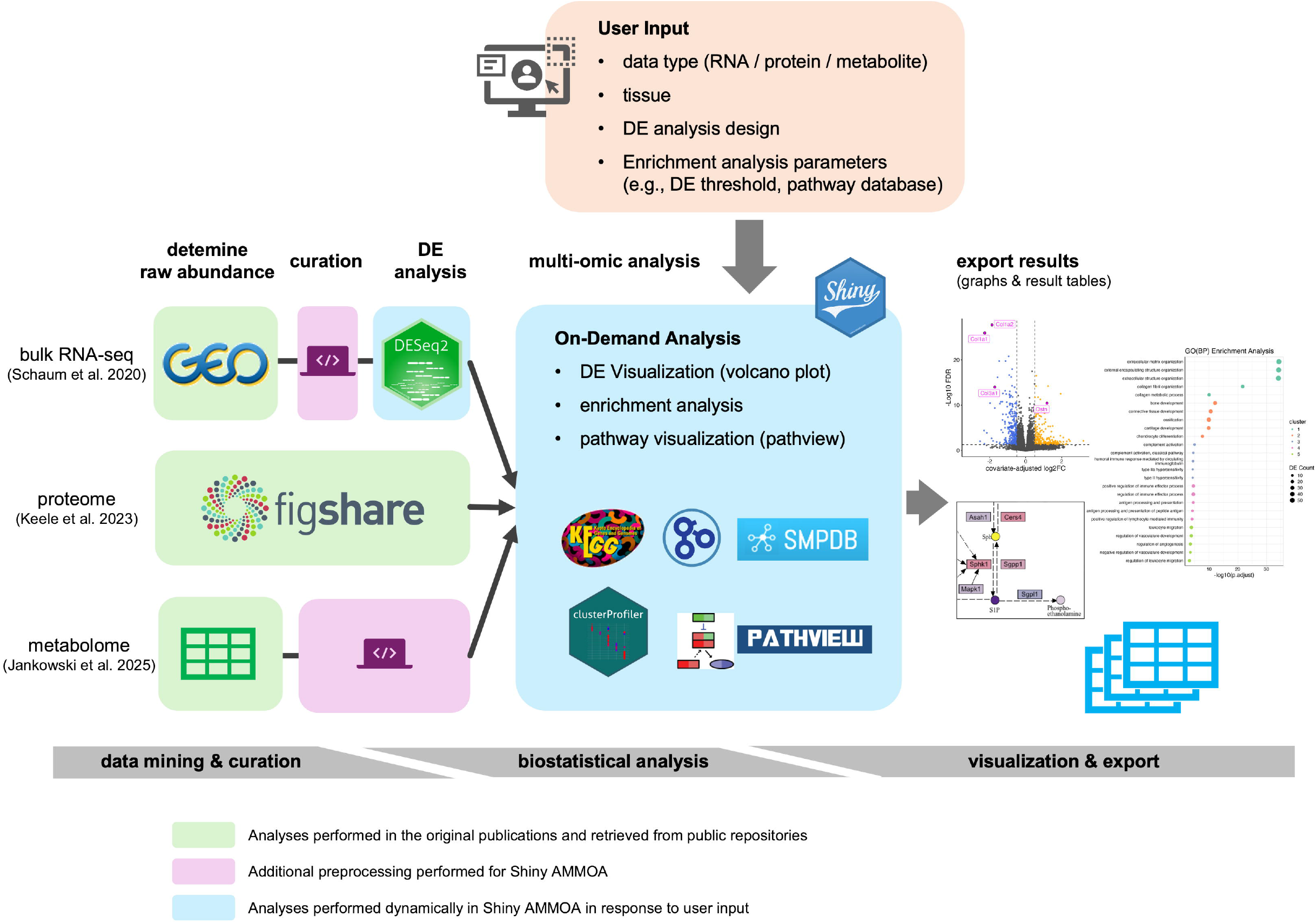
Workflow of Shiny AMMOA implementation. Three omics datasets (bulk RNA-seq, proteomics, and metabolomics) were retrieved from public repositories (see Methods) in either raw or curated form, followed by additional data curation and differential expression analysis when required. Downstream analyses, including pathway enrichment, pathway visualization, and graph rendering, are performed dynamically in response to user input in Shiny application. Analytical results can be downloaded as figures (PNG format) and result tables (CSV format). DE, differential expression.

### Lightweight Web Version Development

The web version of Shiny AMMOA is deployed on the shinyapps.io platform (Posit Software, PBC; https://www.shinyapps.io/). Due to external database terms and conditions, as well as resource limitations such as server memory and CPU capacity of shinyapps.io, several features available in the local version are restricted in the web version. To ensure compliance with external database policies, KEGG pathway diagrams are not displayed because of licensing restrictions (https://www.kegg.jp/kegg/legal.html, last accessed April 24, 2026). This reflects the distinction between service-based provision of KEGG-derived content (web version) and user-initiated access in a local environment (local version), which are subject to different licensing conditions. In addition to these licensing constraints, there are two major differences between the web and local implementations arising from resource limitations. First, for transcriptomic and proteomic pathway enrichment analyses, only KEGG database are supported in the web version. Other databases, such as GO, are available only in the local version, as GO contains a substantially larger number of annotated genes and pathways and therefore requires greater memory resources. Second, the bulk RNA-seq comparisons available for differential expression analysis in the web version are limited to 3-month versus 21-month or all-group analyses, without support for other age-group combinations. This limitation arises because, in contrast to the local version, which performs DESeq2 analysis dynamically in response to user input, the web version visualizes only pre-computed DESeq2 results. Other minor limitation includes that, in transcriptomic analyses, uncharacterized genes (e.g., genes annotated solely with identifiers from the RIKEN Mouse Gene Encyclopaedia Project [19] without corresponding official gene symbols) cannot be displayed.

### Software

All omics data preprocessing was performed in R (version 4.2.2). The local version of Shiny AMMOA was implemented in R (version ≥ 4.2.2) using RStudio (version ≥ 2026.01.0+392) and the Shiny package (version ≥ 1.10.0). Package names and version information for individual analyses are provided in the corresponding sections above.

## Results

### Architecture of the App

Shiny AMMOA is designed to facilitate interactive analysis and visualization of age-associated molecular changes in murine tissues across multiple omics layers. A simplified workflow of Shiny AMMOA development is illustrated in Figure 1. The application supports analyses across three omics layers: bulk transcriptomics, proteomics, and metabolomics. Source datasets for each omics layer were retrieved from publicly available repositories (see Methods). As of version 1.0, Shiny AMMOA consists of five tabs (Figure 2, screen captures): “RNA/Protein Enrichment,” “Metabolite Enrichment,” “Pathway Mapping,” “User Guide,” and “About.” In the “RNA/Protein Enrichment” and “Metabolite Enrichment” tabs, users can perform differential expression analysis and pathway enrichment analysis for individual omics layers. In contrast, the “Pathway Mapping” tab enables users to generate heatmap-colored KEGG pathway diagrams that simultaneously display molecular changes across multiple omics layers.

**Figure 2.**
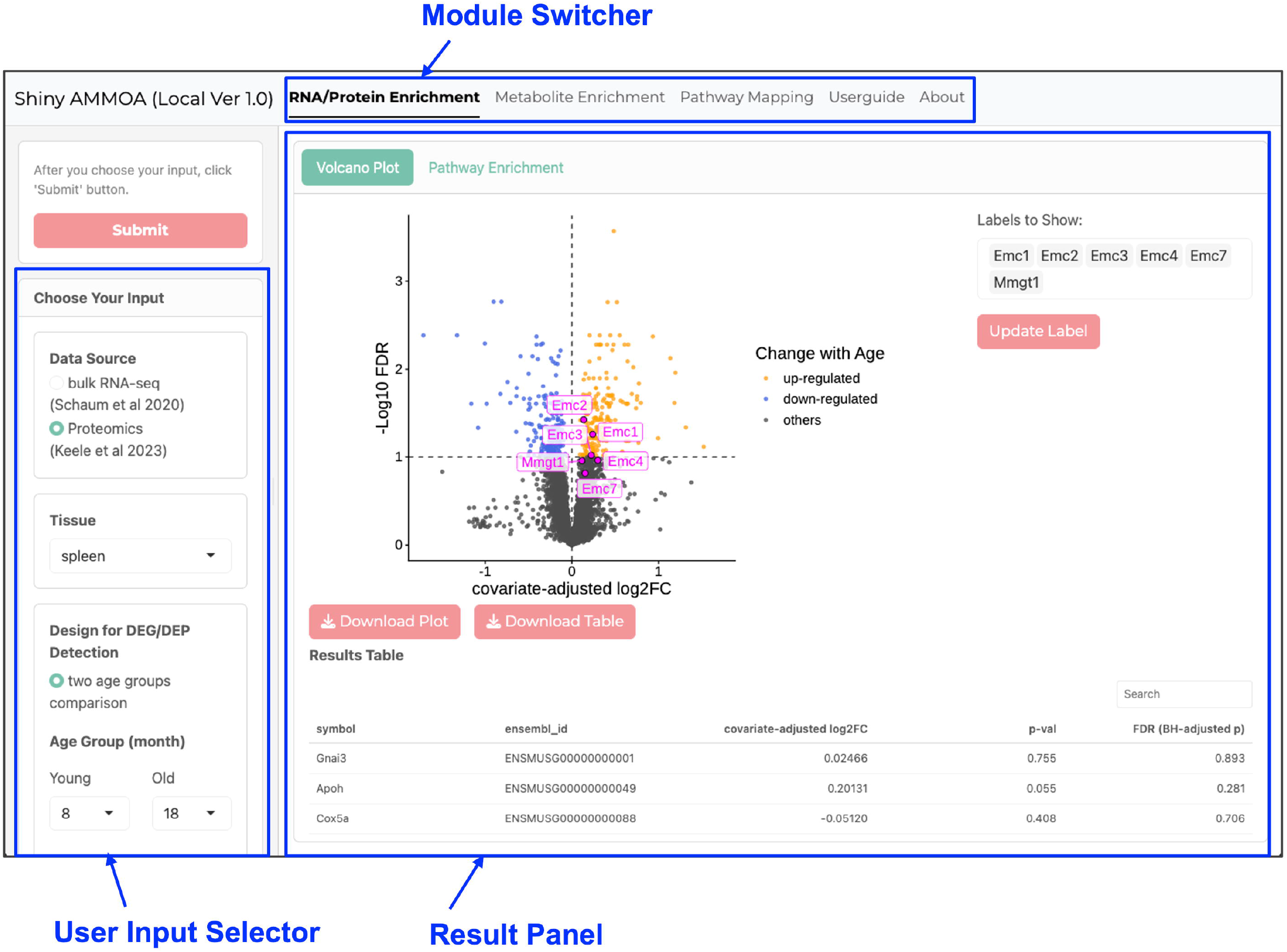
User interface of Shiny AMMOA. Upper panel: module switcher for selecting analytical modules or accessing the user guide. Left panel: user input controls. Main panel: display area for analytical results.

Although Shiny AMMOA is primarily designed to run in a local environment (i.e., on the user’s own computer), a web-based version with limited functionality is available for demonstration purposes at https://m-ninomiya-kanda.shinyapps.io/shiny_ammoa_web/. Details of the system architecture and functional limitations of the web-based version are described in the Methods section.

### Use Case 1: Differential Expression Analysis

A representative output of the “RNA/Protein Enrichment” module is shown in Figure 3a and 3b. To assess reproducibility relative to the original source study, age-associated changes in the spleen proteome were visualized (Figure 3a), followed by Gene Ontology Biological Process (GO(BP)) enrichment analysis of proteins increased with age (Figure 3b). Shiny AMMOA recapitulated key findings of the original publication, including an age-associated increase in endoplasmic reticulum (ER)-related proteins in the spleen [16]. Minor differences between the original publication and Shiny AMMOA outputs are likely attributable to differences in analytical settings, such as pathway redundancy handling and gene set size thresholds used in enrichment testing.

**Figure 3.**
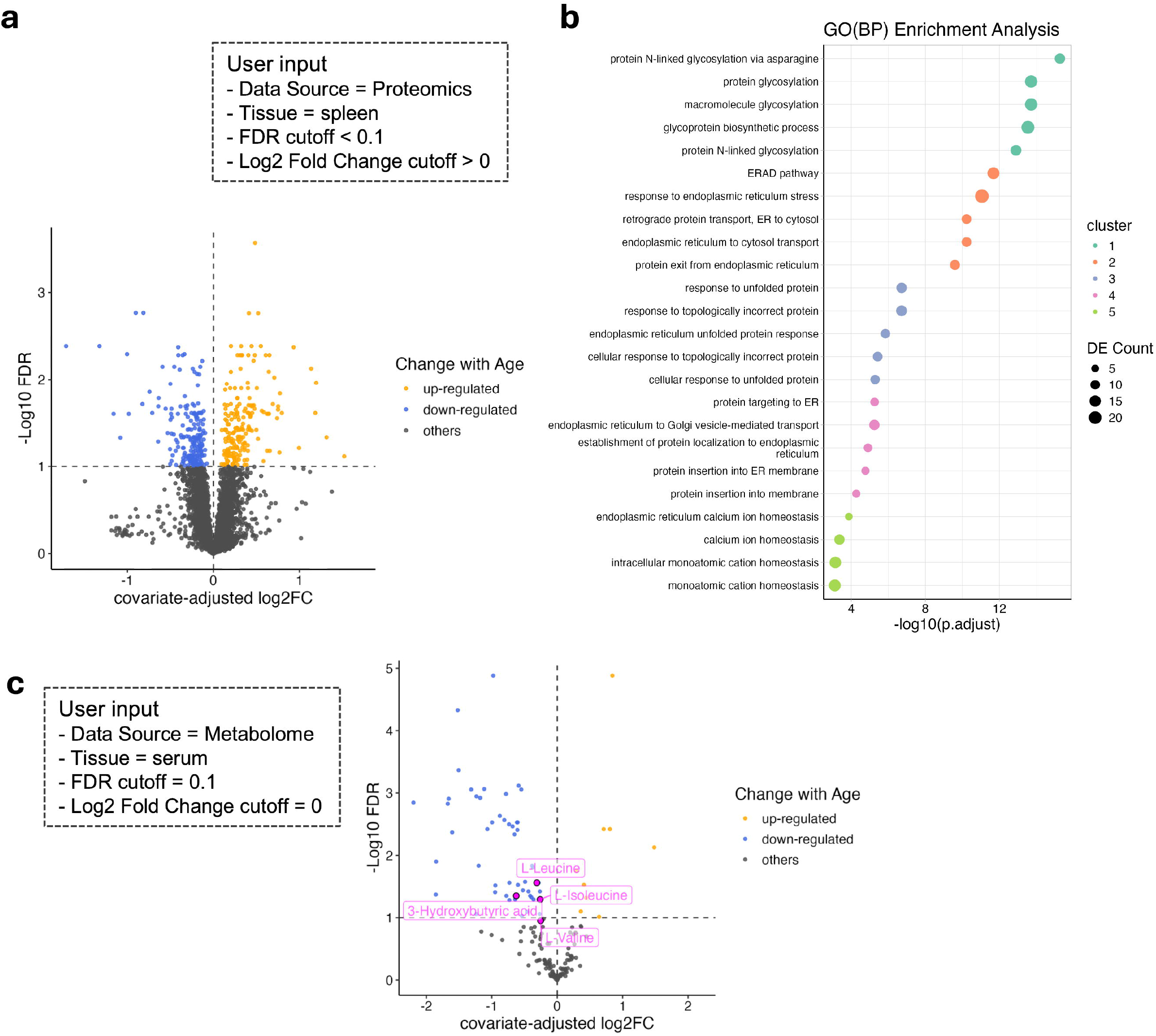
Representative outputs of the enrichment analysis modules in Shiny AMMOA. **(a)** Volcano plot showing age-associated proteomic changes in murine spleen (8 months vs. 18 months). Proteins exceeding user-defined differential expression thresholds (FDR < 0.1, |log2 fold change| > 0) are highlighted. **(b)** Gene Ontology Biological Process (GO(BP)) enrichment results derived from proteins increased with age in (A). Significantly enriched biological terms are clustered based on gene-set similarity, and the top five terms from each cluster are displayed (see Methods). **(c)** Volcano plot showing age-associated changes in circulating metabolites in serum samples (3–6 months vs. >21 months). Metabolites exceeding user-defined differential expression thresholds (FDR < 0.1, |log2 fold change| > 0) are highlighted.

To further illustrate the utility of this feature, a representative result of the “Metabolite Enrichment” module is shown in Figure 3c. Consistent with the original publication [7] and other independent prior studies [20,21], serum levels of 3-hydroxybutyrate, a major circulating ketone body, were significantly decreased with age. Several amino acids, including the branched-chain amino acids L-leucine and L-isoleucine, also showed age-associated decreases as described in the source publication, whereas L-valine exhibited a similar trend but with FDR ≥ 0.1. Notably, tissue-level analyses revealed that 3-hydroxybutyrate was reduced with age in liver and quadriceps muscle but not in soleus muscle (Supplemental Figure 1), suggesting age-associated impairment in hepatic ketone body production and reduced ketone body availability in glycolytic muscle, whereas oxidative muscle appears relatively preserved.

### Use Case 2: Pathway Diagram Visualization

Figure 4a visualizes age-associated expression changes of genes involved in the “ECM–receptor interaction” pathway (KEGG ID: mmu04512) in limb muscle (tibialis anterior). Consistent with the source data article [6] and other independent studies [22–24], many pathway genes exhibited significant age-dependent alterations, potentially reflecting dysregulated extracellular matrix (ECM) remodeling during aging [22]. Importantly, visualization of individual pathway components on the KEGG diagram revealed that expression changes were not uniform across functional categories: while most ECM structural components, such as collagens, showed a monotonic decline with age, integrin genes displayed more subtype-specific patterns. This distinction is biologically informative, as integrins primarily function as ECM receptors rather than structural components [25], highlighting how pathway visualization can reveal functionally heterogeneous aging patterns within a single pathway.

**Figure 4.**
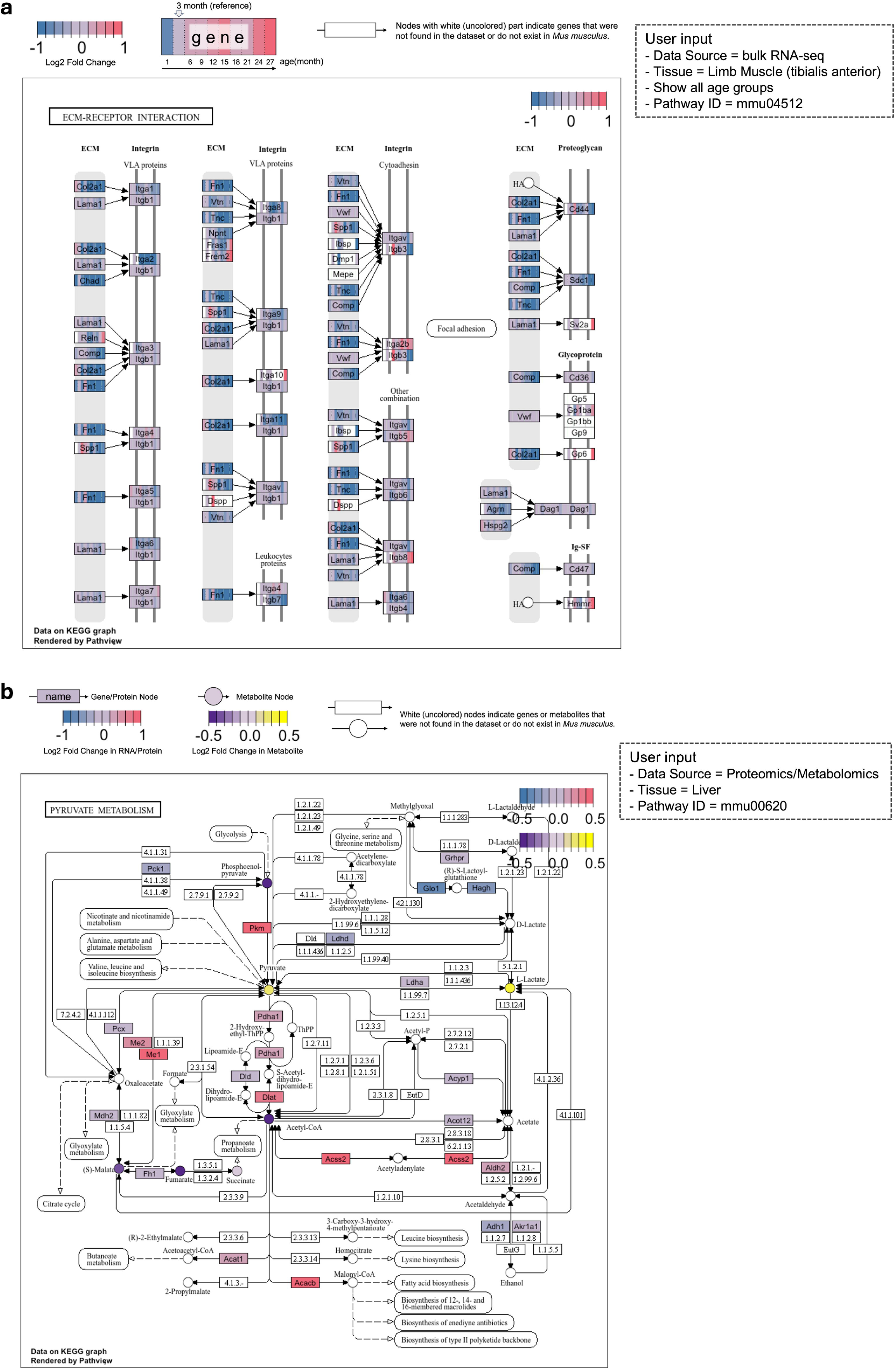
Representative outputs of the pathway mapping module in Shiny AMMOA. **(a)** Age-associated bulk transcriptomic changes in murine limb muscle (tibialis anterior) mapped onto the “ECM–receptor interaction” pathway (KEGG ID: mmu04512). Expression changes across the murine lifespan (1, 3, 6, 9, 12, 15, 18, 21, and 24 months) are visualized using a heat-scale, with the 3-month time point as the reference. **(b)** Age-associated proteomic (8 months vs. 18 months) and metabolomic (3–6 months vs. >21 months) changes in murine liver overlaid onto the “Pyruvate metabolism” pathway (KEGG ID: mmu00620). Rectangular nodes represent protein expression, and circular nodes represent metabolite abundance. Both the KEGG pathway diagram and graphical legend were generated and downloaded directly from Shiny AMMOA.

Finally, to demonstrate how Shiny AMMOA integrates multiple omics layers, age-associated proteomic and metabolomic changes in the “Pyruvate metabolism” pathway (KEGG mmu00620) in liver are visualized in Figure 4b. Pyruvate metabolism occupies a central position in cellular energy homeostasis, linking glycolysis, gluconeogenesis, and the tricarboxylic acid cycle, and plays a critical role in hepatic metabolic flexibility [26], which is known to decline with aging [27]. Although the metabolomic datasets incorporated into Shiny AMMOA do not comprehensively cover all metabolites in this pathway, due to not only limited panel size but detection challenges for certain intermediate metabolites on the LC-MS platform, the visualization remains informative. It enables intuitive assessment of coordinated age-associated changes in metabolic enzymes and measurable metabolites.

## Discussion

In the present report, I developed Shiny AMMOA, a GUI-based, user-friendly analytical platform for integrative analysis of murine aging transcriptomic, proteomic, and metabolomic datasets. By systematically curating publicly available multi-omics resources and implementing a modular Shiny-based architecture, Shiny AMMOA enables end-to-end exploratory analyses, including differential expression, enrichment testing, and pathway-level visualization across individual and multiple omics layers within a unified interface. Through representative use cases, I demonstrated that the application not only recapitulates key findings from original source studies but also facilitates intuitive discovery of tissue-, pathway-, and modality-specific aging signatures. Such features make Shiny AMMOA particularly valuable for experimental biologists and aging researchers who may not be fully familiar with programming, as it enables them to directly interrogate large-scale public datasets, rapidly generate biologically grounded hypotheses, and prioritize pathways or molecular targets for downstream experimental validation without requiring custom scripting or specialized computational infrastructure.

While Shiny AMMOA offers user-friendly, interactive analyses of multi-omics data, certain design choices reflect deliberate trade-offs to ensure accessibility and usability for experimental scientists. First, for transcriptomic analyses, Shiny AMMOA currently incorporates only bulk RNA-seq data. While single-cell RNA-seq (scRNA-seq) and single-nucleus RNA-seq (snRNA-seq) are increasingly recognized as more informative for capturing cell type-specific gene expression changes, bulk RNA-seq aggregates heterogeneous cell populations within tissues, which may attenuate differential expression signals in specific cell types. Indeed, the Tabula Muris Consortium published both age-related murine single-cell and bulk RNA-seq datasets on the same day [6,28]. Nevertheless, Shiny AMMOA adopts bulk RNA-seq data in accordance with its design philosophy; Analyses of single-cell-scale datasets typically involve large data volumes and substantial computational demands, making interactive, GUI-based statistical analyses impractical on entry-level, home-use computers. Therefore, I chose to leverage bulk RNA-seq data, which are sufficiently informative for initial exploratory analyses while minimizing computational burden.

Second, for all three omics layers, pathway enrichment analysis in Shiny AMMOA is performed exclusively using over-representation analysis (ORA). Although ORA is a canonical and widely used method, alternative approaches such as gene set enrichment analysis (GSEA) have become increasingly popular [12]. GSEA evaluates coordinated changes in pre-ordered gene sets without requiring arbitrary thresholds for differential expression and can often achieve higher sensitivity while maintaining acceptable specificity. Despite these advantages, ORA was selected for two main reasons. First, GSEA typically requires greater computational resources, which limits its suitability for dynamic analyses within a GUI-based application. Second, the interpretation of GSEA results may be less intuitive for experimental scientists, as key altered biomolecules are not explicitly defined by differential expression thresholds, such as fold change and FDR, in contrast to ORA. Nevertheless, GSEA and other analysis or visualization methods could be added in future versions of Shiny AMMOA, allowing the application to accommodate additional analytical and exploratory needs in response to users’ feedback.

Beyond these functional trade-offs, Shiny AMMOA also exhibits limitations inherent to the available data and the current implementation. First, the application relies exclusively on publicly available datasets, which may include batch effects or other biases derived from analytical platform that are beyond the control of either the user or the developer. Second, although Shiny AMMOA enables integrative interpretation across multi-omic layers by simultaneously visualizing gene/protein and metabolite changes on the same KEGG pathway diagrams, the underlying differential expression analyses are conducted separately for each omics layer. More comprehensive integration at the data analysis stage, rather than at the level of downstream visualization, therefore remains an important area for future development.

In conclusion, Shiny AMMOA provides an accessible and extensible platform for exploratory, integrative analysis of murine aging-associated multi-omics data, bridging the gap between large-scale public resources and hypothesis-driven experimental research. By lowering computational barriers while preserving analytical rigor, the application enables a broader community of aging researchers to extract biological insights from complex datasets and to more effectively translate systems-level observations into testable experimental questions.

## Data Availability

All source data underlying this work are available in previously published works (see Methods). Curated datasets generated during the current work are available at https://github.com/M-Ninomiya-Kanda/Shiny_AMMOA_preprocessing.

## Code Availability

All code associated with this work, including R scripts for data curation and the source code for Shiny AMMOA, is available at the author’s GitHub repository along with setup instructions. The R scripts for data curation: https://github.com/M-Ninomiya-Kanda/Shiny_AMMOA_preprocessing, source code for Shny AMMOA local version with setup instructions: https://github.com/M-Ninomiya-Kanda/Shiny_AMMOA_local and source code for lightweight web version: https://github.com/M-Ninomiya-Kanda/Shiny_AMMOA_web.

## Author Contributions

Conceptualization, Data curation, Formal analysis, Software, Visualization, Writing – original draft: MNK.

## Acknowledgement

The author would like to thank Irina M. Conboy and Michael J. Conboy for the discussion in preliminary computerization and Masahiro Ninomiya for technical suggestions about software implementation.

## Source of Funding

This work was supported by funding provided by Ajinomoto Co., Inc. to MNK as part of a trainee program. The funder had no role in the study design, data collection, analysis, or interpretation.

## Statements and Declarations

MNK is a full-time employee of Ajinomoto Co., Inc. and is currently affiliated with University of California, Berkeley as part of a trainee program. The employer had no role in the study design, data collection, analysis, or interpretation.

