## Supplemental Figure for "Shiny AMMOA: an interactive platform for integrative multi-omics analysis of murine aging"

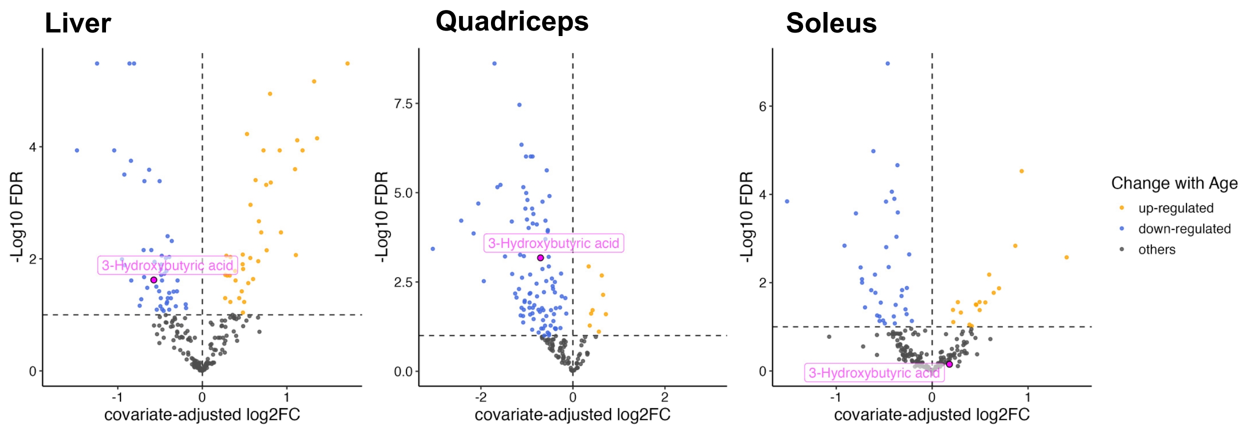


**Supplemental Figure 1. Examples of metabolomic differential abundance analysis**
Volcano plots showing age-associated changes in metabolites from liver, quadriceps, and soleus samples (3–6 months vs. >21 months). Metabolites that exceed user-defined differential expression thresholds (FDR < 0.1, $|\log_2$ fold change$| > 0$) are colored. 3-Hydroxybutyric acid is highlighted using the built-in labeling feature of Shiny AMMOA.
